# Inferring relevant cell types for complex traits using single-cell gene expression

**DOI:** 10.1101/136283

**Authors:** Diego Calderon, Anand Bhaskar, David A. Knowles, David Golan, Towfique Raj, Audrey Q. Fu, Jonathan K. Pritchard

## Abstract

Previous studies have prioritized trait-relevant cell types by looking for an enrichment of GWAS signal within functional regions. However, these studies are limited in cell resolution by the lack of functional annotations from difficult-to-characterize or rare cell populations. Measurement of single-cell gene expression has become a popular method for characterizing novel cell types, and yet, hardly any work exists linking single-cell RNA-seq to phenotypes of interest. To address this deficiency, we present RolyPoly, a regression-based polygenic model that can prioritize trait-relevant cell types and genes from GWAS summary statistics and single-cell RNA-seq. We demonstrate RolyPoly’s accuracy through simulation and validate previously known tissue-trait associations. We discover a significant association between microglia and late-onset Alzheimer’s disease, and an association between oligodendrocytes and replicating fetal cortical cells with schizophrenia. Additionally, RolyPoly computes a trait-relevance score for each gene which reflects the importance of expression specific to a cell type. We found that differentially expressed genes in the prefrontal cortex of Alzheimer’s patients were significantly enriched for highly ranked genes by RolyPoly gene scores. Overall, our method represents a powerful framework for understanding the effect of common variants on cell types contributing to complex traits.

## Introduction

Identifying the primary subset of cell types or states and genes involved in complex traits is critical to the process of developing mechanistic insights. For example, knowledge that the *FTO* locus acts on *IRX3* and *IRX5* primarily in human adipocyte progenitor cells enabled researchers to rigorously define a novel thermogenesis pathway central for lipid storage and obesity [6]. And, focusing on distinct human *C4* isotypes, Sekar et. al., highlighted the role of the classical complement cascade (of which *C4* is a critical component) and synapse elimination during development in the brains of individuals with schizophrenia [57].

In addition to estimating disease risk for individual variants, GWAS have proven useful for identifying trait-relevant cell types or tissues. Assuming variants affect phenotypes through gene regulation, one can prioritize cell types for further analysis with an enrichment of GWAS signal in cell type-specific functional regions of the genome that affect gene regulation. A series of studies identified enrichment of GWAS signal in sorted cell type [51] or tissue-specific eQTLs [45]. Other approaches have revealed enrichment of GWAS signal in cell type-specific functional annotations (e.g., ATAC-seq, ChIP-seq, RNA-seq) [25, 66, 47, 15, 61, 14, 12]. However, these analyses are limited in cell type resolution because they either require samples with population variation (infeasible to collect for many cell types) or rely on functional assays that require on the order of thousands of cells, which are challenging to collect for rare or uncharacterized cell types. Thus, it remains difficult to evaluate whether disease phenotypes are driven by tissues, broad cell populations, or very specific cell types. Furthermore, an inability to analyze difficult-to-characterize cell types is a concern when scanning for links between traits and cell types in complex tissues composed of many heterogenous cell types. For example, describing the brain as the primary pathogenic tissue responsible for schizophrenia or Alzheimer’s disease is unsatisfying, but it remains difficult to comprehensively collect functional information from the plethora of brain cell types necessary to perform standard GWAS enrichment analyses.

Meanwhile, single-cell gene expression technology has offered insights into complex cell types [46, 27, 71, 65, 33, 16, 20, 28, 4]. Additionally, there are concerted efforts underway to develop comprehensive single-cell atlases of complex human tissues known to be associated with phenotypes of interest, such as immune cell types for autoimmune disease and brain cell types for neuropsychiatric disorders [52]. However, to our knowledge, there are no existing methods designed to link novel single-cell based cell types and phenotypes of interest.

Thus, we developed RolyPoly, a model for prioritizing trait-relevant cell types observed from single-cell gene expression assays. Importantly, RolyPoly takes advantage of polygenic signal by utilizing genome-wide GWAS summary statistics for all SNPs near protein coding genes, appropriately accounts for linkage disequilibrium (LD), and jointly analyzes gene expression from many tissues or cell types simultaneously. Additionally, our model can utilize signatures of cell-specific gene expression to prioritize trait-relevant genes. Finally, we provide a fast and publicly available implementation of the RolyPoly model.

## Material and Methods

### Overview of the methods

The primary goals of RolyPoly are to identify and prioritize trait-relevant cell types (or tissues) and genes (Figure 1). At a high-level, RolyPoly starts by learning about the relationship between gene expression and estimated GWAS effect sizes from a trait of interest (captured with our *γ* model parameters, described below). For example, we might expect to observe larger GWAS effect sizes for cholesterol regulation at SNPs that affect liver-specific gene expression because the liver is known to regulate cholesterol levels. Thus, based on such an enrichment, RolyPoly would learn that the liver is a trait-relevant tissue. Next, we can use this knowledge to prioritize trait-relevant genes by calculating a score (represented by 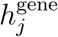, defined below) that identifies genes upregulated in RolyPoly-inferred relevant tissues. Continuing with our example, once we know that liver-specific gene expression is associated with larger GWAS effect sizes, RolyPoly would prioritize studying liver-specific genes in the context of understanding cholesterol regulation (resulting in larger 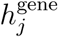 values). Below we describe the details of how RolyPoly carries out each of these steps.

**Figure 1:**
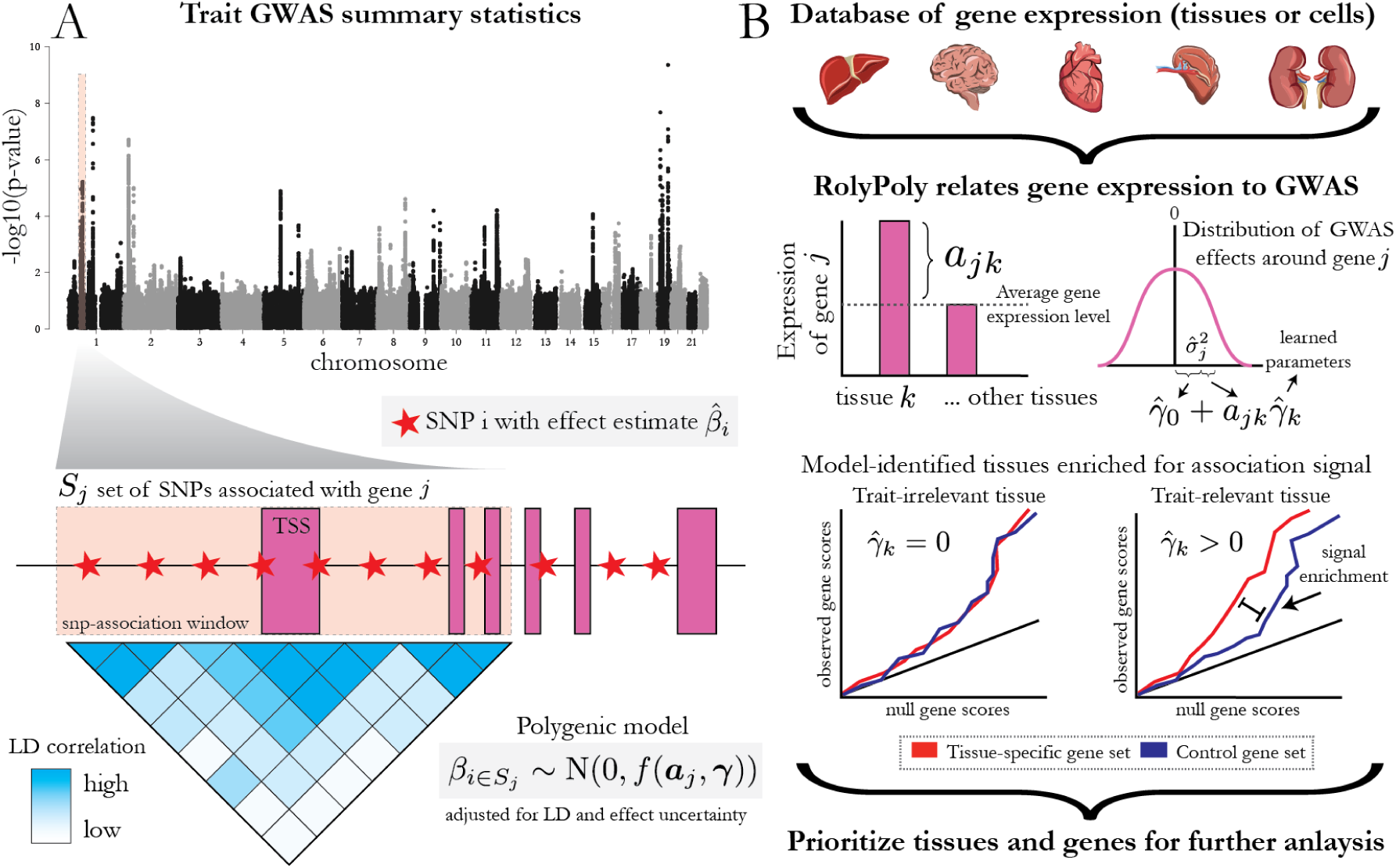
RolyPoly detects trait-associated annotations using GWAS summary statistics and gene expression profiles. A) We model the variance of GWAS effect sizes of SNPs associated with a gene as a function of gene annotations, in particular gene expression, while accounting for LD using population matched genotype correlation information. (Manhattan plot is based on data from [19].) From a database of functional information (such as tissue or cell type RNA-seq) we learn a regression coefficient for each annotation, 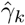, that captures its influence on the variance of GWAS effect sizes. A deviation from the mean gene expression value of a_jk_ results in an increase of 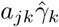 to the expected variance of gene-associated GWAS effect sizes. The value 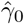 represents a regression intercept that estimates the population mean variance. To check learned model parameters, we expect to see an enrichment of LD-informed GWAS gene scores for genes that are specifically expressed in a tissue inferred to be trait relevant. Finally, from a model fit we can prioritize trait-relevant tissues and genes.

### GWAS summary statistics

Consider a fully polygenic GWAS model 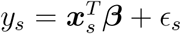, where *y*_*s*_ is the phenotypic measurement from individual *s*, ***x***_*s*_ is a vector of genotypes at *p* SNPs for individual *s*, ***β*** is a vector of *p* SNP effects, and we represent the stochastic environmental error with 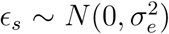. Importantly, we assume that the matrix of genotypes has been scaled and standardized such that the mean is 0 and variance 1 for each SNP vector (and similarly for the trait *y*_*s*_). The main summary statistics released by GWAS are per-variant effect estimates, which we refer to as 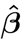. Researchers typically calculate and report univariate effect-size estimates. These estimates represent the marginal standardized regression coefficient and are calculated as 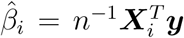, where ***X***_*i*_ (note the change in case) represents standardized genotypes for SNP *i* across the *n* individuals (see Appendix for derivation). Substituting the polygenic model for ***y*** into the estimation equation (see Appendix for derivation), the sampling distribution of the estimated SNP effect sizes corresponds to

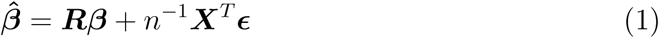

where ***R*** is the sample LD matrix (i.e. *r*_*ii′*_ is the Pearson correlation values between genotype *i* and *i′*). Using this definition of estimated GWAS effect sizes, we develop a highly polygenic approach that models the variance of these SNP effect sizes as a function of annotation specificity of proximal gene expression.

### Polygenic model

For notational convenience, let *g*(*i*) represent the gene associated with SNP *i* and *S*_*j*_ = {*i*: *g*(*i*) = *j*} be the set of SNPs associated with gene *j*. We use the notation ***β***_*S*_ to denote the coordinates of ***β*** whose indices lie in set *S*. We assume *a priori* that the true GWAS effect sizes of SNPs in gene *j* follow a normal distribution *β*_*S*_j__ ∼ MVN(0, τ_*j*_*I*), where *I* is the *|S*_*j*_*|* × *|S*_*j*_*|* identity matrix, and *τ*_*j*_ is the prior effect size variance for all SNPs associated with gene *j* and is modeled as a linear function. More specifically, *τ*_*j*_ is a linear function of *N* annotations *a*_*jk*_ (in this case cell-type specific gene expression), with annotation coefficients *γ*_*k*_ and an intercept term *γ*_0_:

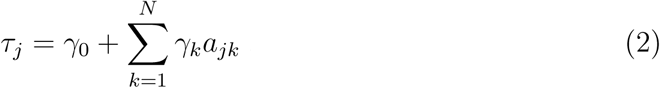

RolyPoly estimates the parameter vector ***γ***, which captures the influence of cell-type specific gene expression on the variance of GWAS effect sizes (see Figure 1B). Intuitively, if we estimate a large coefficient for annotation *k*, then we expect larger GWAS effect sizes around genes that are specifically expressed in annotation *k*. On the other hand, it is possible to estimate negative values for some annotation coefficients *γ*. SNPs proximal to genes that are specifically expressed in an annotation with a negative *γ* estimate are expected to have reduced effect size variance compared with the population mean.

Based on this polygenic model, the expected value of the vector of GWAS effect sizes around gene *j* is 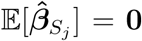, and the covariance matrix is given by 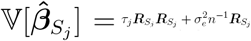, where ***R***_*S*_*j*__ denotes the principal submatrix of ***R*** indexed by the SNPs in *S*_*j*_ (see Appendix for derivation). This model assumes that the effect size of each SNP around a gene *j* is drawn from a distribution with a mean of zero and the same per-SNP variance of *τ*_*j*_. However, there are other SNP annotations that we expect to affect the variance of a GWAS effect size, such as the minor allele frequency (MAF) of the SNP. Therefore, we include *P* SNP-level features as covariates while estimating the variance contribution of gene expression. Specifically, we modify our model to use a per-SNP variance *v*_*i*_ for SNP *i*, given by

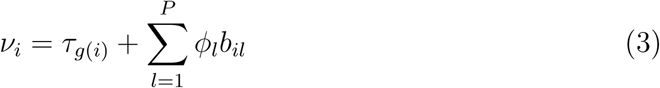

where *τ*_*g*(*i*)_ is the previously described (equation 2) contribution of gene-level annotations to the variance of SNP *i*, *b*_*il*_ is the *i*-th value of SNP-level annotation *l* for SNP *i*, and *φ*_*l*_ is the annotation coefficient for annotation *l*. The distribution for the vector of SNP effects associated with a gene becomes

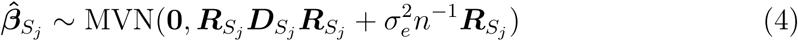

where ***D*** = diag(***v***) is a diagonal matrix of SNP effect size variances. With this modification, we can estimate gene annotation regression coefficients while controlling for the contribution of SNP annotations to the variance of a SNP effect size. We present inferred parameter estimates including accounting for MAF as a SNP-level covariate. MAF values were downloaded from matched population samples from the Phase 3 VCFs of 1000 Genomes Project [1].

For results presented here, we used a window size of 10kb centered on the transcription start site of a gene to associate a SNP to a gene. We chose this window-size because previous work has found that, across a diverse set of cell types and tissues, most eQTLs consistently lie in this region [9, 17, 62, 41]. However, the model description as presented generalizes to larger window sizes or alternative approaches of SNP-gene association. One could rely on enhancer or chromatin maps from ENCODE to incorporate potentially functional variants that are farther away from the TSS. However, doing so would bias our analysis towards well-characterized cell types; thus, we did not include distal elements. With this definition of SNP-gene association there are a few SNPs with multiple associated genes. We duplicate these SNPs and treat them as independent SNP-gene pairs. Since RolyPoly infers parameters from hundreds of thousands of SNPs, we do not expect this contribute significantly to inferred parameters.

### Parameter inference

In order to perform maximum likelihood inference under our model, we would have to compute the determinant and inverse of the potentially high dimensional covariance matrices involved in (4), which would be computationally challenging. Instead, we adopt a method of moments approach, where we fit the gene-level annotation coefficients *γ*_*k*_ and, if included, the SNP-level annotation coefficients *φ*_*l*_. If only gene-level annotations are used, we fit the observed and expected sum of squared SNP effect sizes associated with each gene, where the expected value is given by

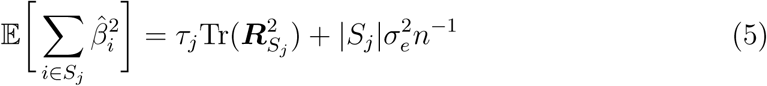

where *Tr* above represents the trace of a matrix (derivation in Appendix). This expectation was derived recognizing that the expected value of the squared *ℓ* _2_ norm of a mean zero multivariate normal distribution is the trace of the covariance matrix. When we include SNP annotation coefficients such that each SNP effect size has a variance term *v*_*i*_, we perform inference by fitting the observed and expected squared effect size of each SNP, where the expected value is given by

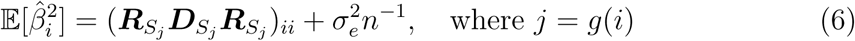

and (***R***_*S*_*i*__ ***D***_*S*_*i*__ ***R***_*S*_*i*__) is the diagonal element of the matrix corresponding to SNP *i*. Interestingly, by using an indicator function rather than quantitative features, we noticed that this model relates to previous work [5] (described in the Appendix). We perform block bootstrap [10] to estimate standard errors, *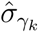*, which are used to compute a *t*-statistic, *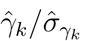*, and corresponding p-values. We use a *t*-statistic because we use our bootstrap estimate of the standard error rather than a known value. The purpose of the block bootstrap is to maintain correlations present in the data when sampling from the empirical distribution, thus, we partitioned the genome into 100 non-overlapping blocks and sample from these blocks with replacement [35]. Additionally, from the bootstrap parameter estimates, we calculate empirical 95% confidence intervals for each 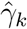. Unless otherwise specified, for our analyses we performed 10^3^ block bootstrap iterations. After including an intercept term, 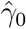, we rank tissues by strength of association with the *t*-statistic or corresponding *p*-value. As in standard regression, the intercept term estimates the population mean of the response term, which in this case is the per-SNP variance of a GWAS effect size.

### Computing trait-relevance gene importance scores and proportion of variance explained by individual annotations

Using a set of inferred gene annotation coefficients, **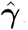**, we calculate several quantities that summarize the contributions of gene annotations to the phenotypic variance. First, we compute 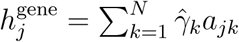, which can be used to rank trait-relevant genes. Essentially, 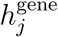 is a gene expression-based prediction of the variance parameter for gene *j* of a normal distribution from which *cis*-GWAS effect sizes are drawn (Figure 1B). Thus, if 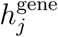 is large we would expect larger *cis*-GWAS effect sizes. Note that this value does not directly rely on GWAS effect size estimates. Instead, 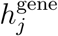 relies on GWAS indirectly through the RolyPoly-inferred parameters. Additionally, where *M* is the number of genes. Through simulation we show that the true value we calculate the contribution of an annotation *k* to a trait as 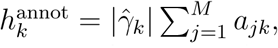, of 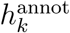 affects our power to detect trait-annotation associations. The total contribution explained by all annotations, *h*^total^, comes from summing the individual annotation values, 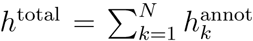. Finally, the proportion of an annotation’s unique contribution to the variance of SNP effects, 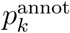, can be calculated as 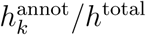.

To validate our gene importance values, 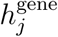, we compare them to gene importance estimates based on *cis*-GWAS summary statistics and LD information. This gene score is an estimate of the variance of GWAS effect sizes accounting for inflation due to local LD, thus we refer to it as the LD-informed gene score. For this calculation we use the methodology described in [38, 37]. However, we use the same window size around a gene as was used for RolyPoly. In addition to validating 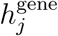, we use the LD-informed gene score to verify GWAS enrichment in specifically expressed genes of model-identified trait-relevant tissues (i.e., Q-Q plots in the Results section).

If the main objective is to compute gene values, 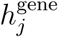, and unbiased parameter estimates are not required, then we include a penalty on the *ℓ*_1_ norm of the annotation coefficients. The penalty strength is modulated with a *λ* tuning factor which is chosen based on cross validation. Regularization has the beneficial effect of shrinking parameter estimates of irrelevant tissues and can result in higher gene score prediction accuracy.

### Simulation setup

For clarity we denote generated parameters and data with an asterisk (*). In simulation results reported we used 2 × 10^4^ genes, five simulated gene annotations and one simulated SNP annotation. We generated gene expression, *a*^*\**^, from a standard *χ*^2^-distribution, and allele frequency as an example SNP annotation, *b*^*\**^, from a standard uniform distribution. Recall that our model annotation coefficients determine the influence these annotations will have on SNP effect sizes. For each simulated data set we fixed annotation effects by sampling from a uniform distribution, *ф*^*\**^ ∼ Uniform(0, 10^−5^) for SNP annotation effects and *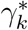* ∼ Uniform(0, 10^−5^) for gene annotation effects. We combined the simulated functional information and annotation coefficients to calculate a per-SNP variance term. Thus, for each SNP effect we computed *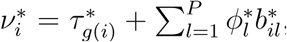*, where 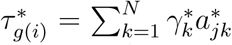. We combined this per-SNP variance term with a per-SNP environmental error contribution set to 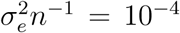 to arrive at the distribution from which we generated simulated effects,

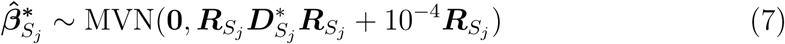

where ***D***^*\**^ is a diagonal matrix with simulated per-SNP variance values. From this distribution, for each simulated gene we sampled 20 SNP effects. As input our inference model takes SNP effects, environmental errors (here set to 10^−4^) and annotations, and attempts to identify the true annotation effects. From this setup we determined whether our method implementation could accurately infer generated SNP annotation effects, 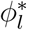, and gene annotation effects 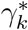.

Although our method assumes each SNP effect size is drawn from the model distribution, it is likely that some GWAS effect sizes come from a null distribution. To test robustness to this potential model misspecification, we first sampled pergene Bernoulli random variables *π*_*j*_ ∼ Bernoulli(*c*), where *c* represents the fraction of causal genes (causal here simply implies sampling from the non-null model). We sampled SNP effects for each gene as

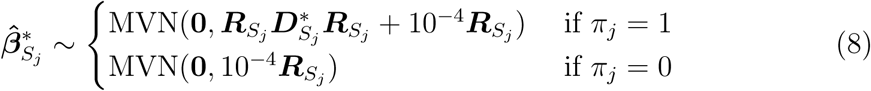

Varying the fraction of causal genes, parameter *c*, across simulated data sets, we studied its effect on model inference.

### Obtaining gene expression databases and GWAS summary statistics

We estimated annotation parameters for three gene expression databases. 1) The Genotype-Tissue Expression (GTEx) cohort includes RNA-seq from different individuals at many tissue sites [39]. 2) We downloaded single-cell RNA sequencing data from Ziesel et al., containing data for 3005 single cells from the hippocampus and cerebral cortex of mice [71]. 3) We obtained human single-cell RNA sequencing data of cortex samples from Darmanis et al., [7]. Within each gene expression database we standardized the distribution of gene expression across samples with quantile normalization. Expression samples from the same tissue or purified cell population were averaged. In the case of single-cell expression data we took the average of single-cell expression vectors for common previously defined cell type classes. To compare across genes, we scale, center, and then square expression values across annotations. When using an expression database from mice, we only used orthologous protein coding genes with a one-to-one functional mapping (based on the definition in Ensembl’s BioMart [31]).

We downloaded publicly available GWAS summary statistics from 10 traits from their respective publications: Schizophrenia [43], late-onset Alzheimer’s disease [36], four metabolic traits from [19] (HDL cholesterol, LDL cholesterol, total cholesterol and Tryglyceride levels), educational attainment [44], height [70], extreme body mass index [3], and age-related cognitive decline [8]. We restricted or analysis to the autosomes, removed the *MHC* region for immune traits (chromosome 6 between 25 and 34 Mb), and removed rarer variants (MAF *<* 0.1%). For late-onset Alzheimer’s disease and age-related cognitive decline, in addition to using the entire set of GWAS summary statistics, we ran RolyPoly after removing variants from a 1 Mb window centered on the TSS of *APOE* (chromosome 19 between 44909011 and 45909011). All referenced genome coordinates are from hg19.

### Differential gene expression analysis

For the analysis of 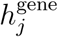 enrichment in differentially expressed genes of Alzheimer’s patients, we downloaded microarray gene expression data from 230 samples of the prefrontal cortex [72]. We used Limma to perform a differential gene expression analysis between patient and control tissues [55]. Probes were mapped to genes using a mapping downloaded from Ensembl’s BioMart [31]. If multiple probes mapped to a single gene we took the median expression value across all probes. Unless otherwise specified, we performed Kolmogorov-Smirnov significance tests of gene value enrichment within differentially expressed genes.

### Calculating RolyPoly gene score enrichment accounting for correlations among gene expression values

To assess the enrichment of RolyPoly gene scores among differentially expressed genes we calculate the Spearman rank correlation coefficent, *ρ*_obs_, between RolyPoly gene scores and a differential expression *t*-statistics. A large value of *ρ*_obs_ indicates enrichment of large RolyPoly gene scores among differentially expressed genes. Assessing the significance of *ρ*_obs_ by considering each gene as independent will be anti-conservative because of correlation between gene expression levels of co-regulated genes. To account for this, we generate an empirical sampling distribution for *ρ* under the null of no association between RolyPoly scores and *t* which accounts for gene expression correlation.

We estimate the variance-covariance matrix of gene expression in healthy individuals, Σ. Because there are fewer samples than genes we use singular value decomposition (SVD) to represent the low-rank Σ matrix. Under the null hypothesis, we generate a gene expression matrix for both case and control samples using the same distribution, ***X***_*i*_ ∼ MVN(0, Σ). We have two sets of individuals, the set of healthy controls, *H*, and the set of affected individuals, *A* (of equal size to the true data). For each gene *j* we compute a *t*-statistic testing the difference between the means of the healthy and affected simulated expression values,

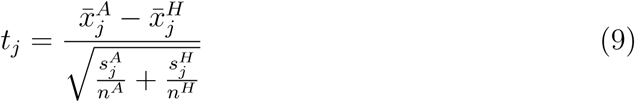

where *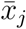* is the mean expression of gene *j*, *s*_*j*_ is the sample variance, and *n* is the sample size. We compute Spearman’s correlation coefficient *ρ*_sim_ between *t*_*j*_ and 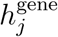. We repeat the process of generating expression and calculating *ρ*_sim_ 10^3^ times to generate a null distribution which is then used evaluate the significance of *ρ*_obs_.

### Calculating LD correlation values

We downloaded Phase 3 VCFs of European individuals from the 1000 Genomes Project [1]. We used PLINK v1.90b1b to calculate Pearson’s r values of SNPs within the default 1 Mb window [49].

### RolyPoly implementation and usage

We implemented our method for use through the RolyPoly R package, which is made available free and open source via CRAN and at our git repository (see Web Resources).

## Results

### Simulation

We used simulations (see Material and Methods) to verify our implementation of RolyPoly and characterize properties of parameter estimation and hypothesis testing.

Across 500 data simulations, we found that RolyPoly-inferred *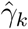* parameters were unbiased estimates of the true underlying effect *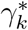* (see Figure 2A). This is an important property if we aim to accurately quantify the total contribution of an annotation to a trait, 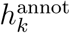. 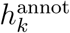 summarizes the amount of signal present in the dataset to detect an association between the trait and annotation *k*. In particular, our power is strongly dependent on 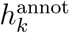 (see Figure 2B), where power refers to the probability that we correctly reject the null hypothesis (i.e., 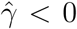). It is likely that some fraction of GWAS effect sizes are drawn from a null distribution, which we do not currently model in RolyPoly. Thus, we investigated the effect of varying the fraction of GWAS effects drawn from the model distribution and our power to detect significant annotations. As expected, when the fraction of genes simulated from the causal distribution decreases we lose power (see Figure 2B). However, even with 25% of genes (and downstream GWAS effect sizes) drawn from the causal distribution, we achieve greater than 50% power for an annotation with 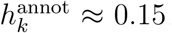. For context, in real data, we consistently observed 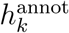 values greater than 0.1.

**Figure 2:**
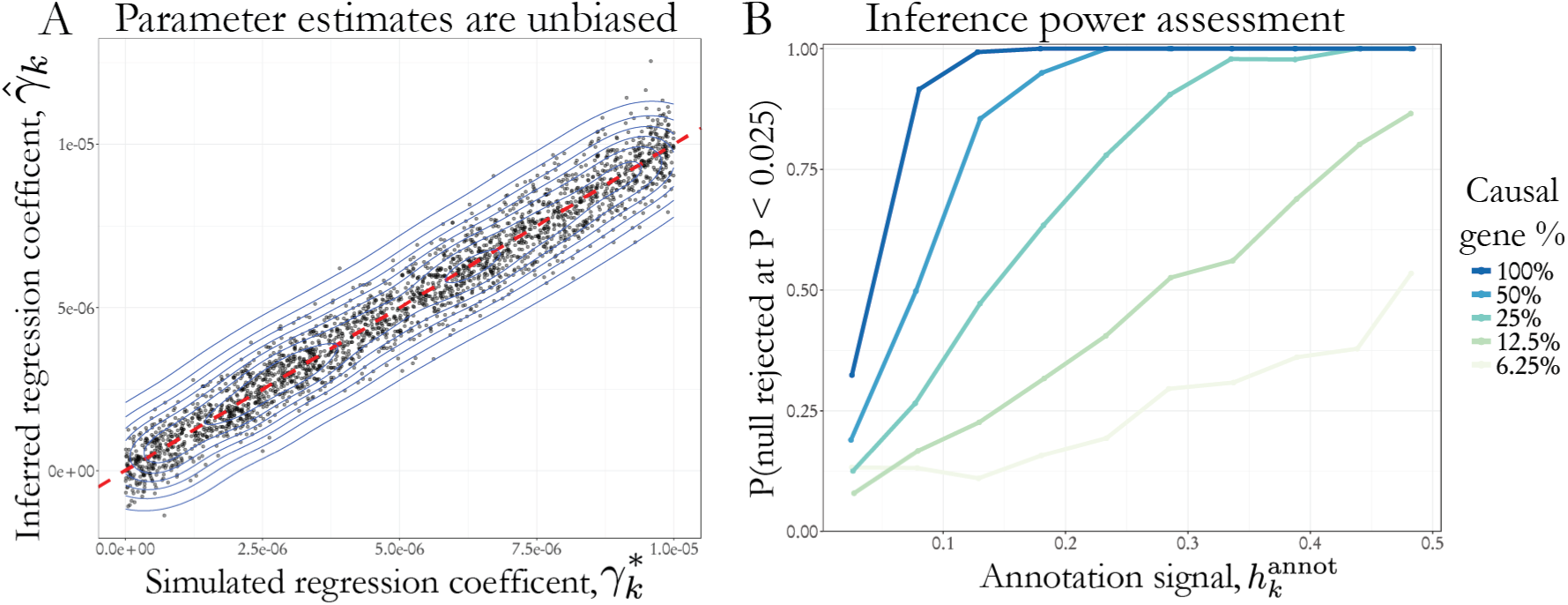
Simulation results. A) Parameter inference is unbiased and accurate for a range of simulated γ^*^ effects. Red-dashed line represents the identity line. B) Power as a function of the 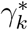 and annotation values defined as 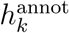 in the Material and Methods section. Even when some SNPs are drawn from the null distribution we maintain reasonable power to detect associations.

For data generated under the model, we demonstrated that our estimated parameters are unbiased and have low levels of deviation around the true parameter values. Our power to detect significant annotations is modulated by the annotation effect, the annotation values, and the fraction of effects drawn from the model distribution. Furthermore, in the setting where the effects are simulated from a mixture of the model and null distribution we still have power to detect significant annotations.

### Trait-relevant tissues identified from GTEx data

As a proof-of-principle, we ran our method on trait-association data from publicly available GWAS traits and gene expression data from 27 tissues of 544 individuals from the GTEx consortium (data download and processing described in Material and Methods).

In Table 1, we summarize the top two tissue-trait associations that pass a marginal significance threshold (*p <* 0.05) for seven GWAS traits. With an extreme body mass index GWAS (BMI) we found associations with kidney (*p* = 7 × 10^−3^) and thyroid (*p* = 0.03) tissue gene expression. Obesity is known to negatively affect kidney function; however, from existing literature it is ambiguous whether the tissue has a causal role in determining body mass index [23]. There are studies that demonstrate a correlation between thyroid function and weight [53, 32]. We observed a significant enrichment of educational attainment (EA) signal for genes specifically expressed in the pituitary gland (*p* = 0.03) and brain (*p* = 0.04), which corresponds with recent analysis [54, 44]. For height, we detect an association with muscle (*p* = 6 × 10^−10^) and pituitary (*p* = 6 × 10^−7^). Interestingly, tumors in the pituitary are known to lead to gigantism characterized by excessive growth and height [11]. Finally, for several metabolic traits (TC, LDL, TG, HDL), there were signals for the liver, small intestine, and adrenal gland, all of which follow known biology.

**Table 1:**
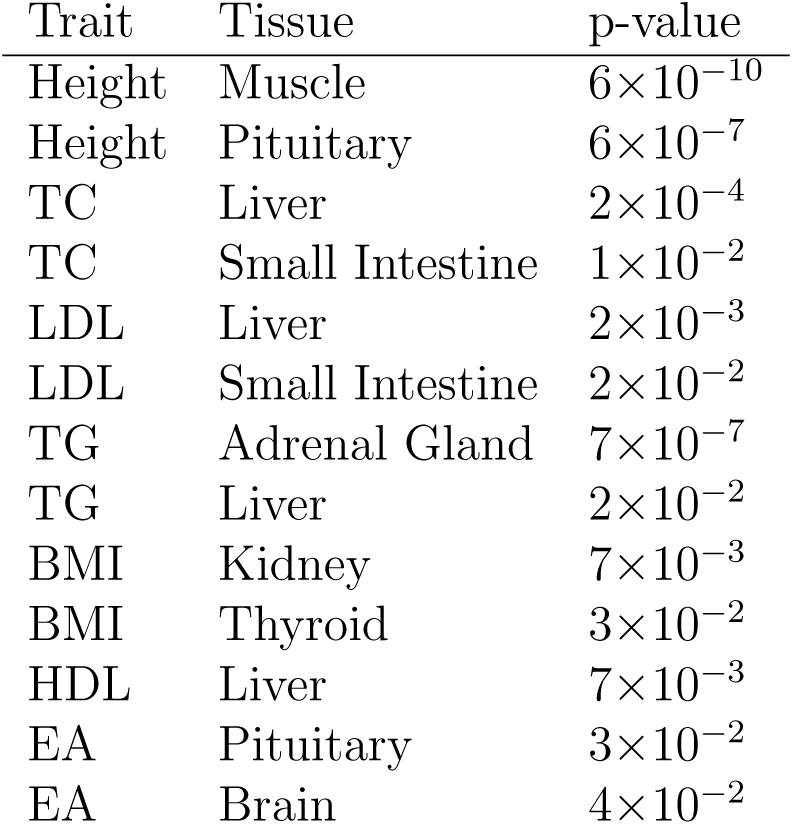
Top trait-relevant GTEx tissue for seven GWAS traits and uncorrected p-values.

| Trait | Tissue | p-value |
| --- | --- | --- |
| Height | Muscle | $6 \times 10^{-10}$ |
| Height | Pituitary | $6 \times 10^{-7}$ |
| TC | Liver | $2 \times 10^{-4}$ |
| TC | Small Intestine | $1 \times 10^{-2}$ |
| LDL | Liver | $2 \times 10^{-3}$ |
| LDL | Small Intestine | $2 \times 10^{-2}$ |
| TG | Adrenal Gland | $7 \times 10^{-7}$ |
| TG | Liver | $2 \times 10^{-2}$ |
| BMI | Kidney | $7 \times 10^{-3}$ |
| BMI | Thyroid | $3 \times 10^{-2}$ |
| HDL | Liver | $7 \times 10^{-3}$ |
| EA | Pituitary | $3 \times 10^{-2}$ |
| EA | Brain | $4 \times 10^{-2}$ |

Next, we examined the total cholesterol (TC) GWAS [19], as its association with liver has been unambiguously reported in the literature. For inference, we used a total of 121,312 SNPs that were within 5kb of a protein coding gene. With *p*-values from our model we ranked tissues by the strength of association with total cholesterol (see left panel of Figure 3). As expected, liver was the clear top-associated annotation (*p* = 2 × 10^−4^), and we estimated an annotation coefficient of 4 × 10^−6^ (see right panel of Figure 3). Thus, we estimated that the variance of TSS-proximal GWAS effect sizes increase by 4 × 10^−6^ as normalized gene expression in the liver increases by one unit (see Material and Methods for a description of gene expression normalization). The small intestine was marginally associated (*p* = 0.01), which follows from the fact that this organ has a central role in nutrient absorption. Additionally, we observed some signal for spleen (*p* = 0.04) and adrenal gland (*p* = 0.05).

**Figure 3:**
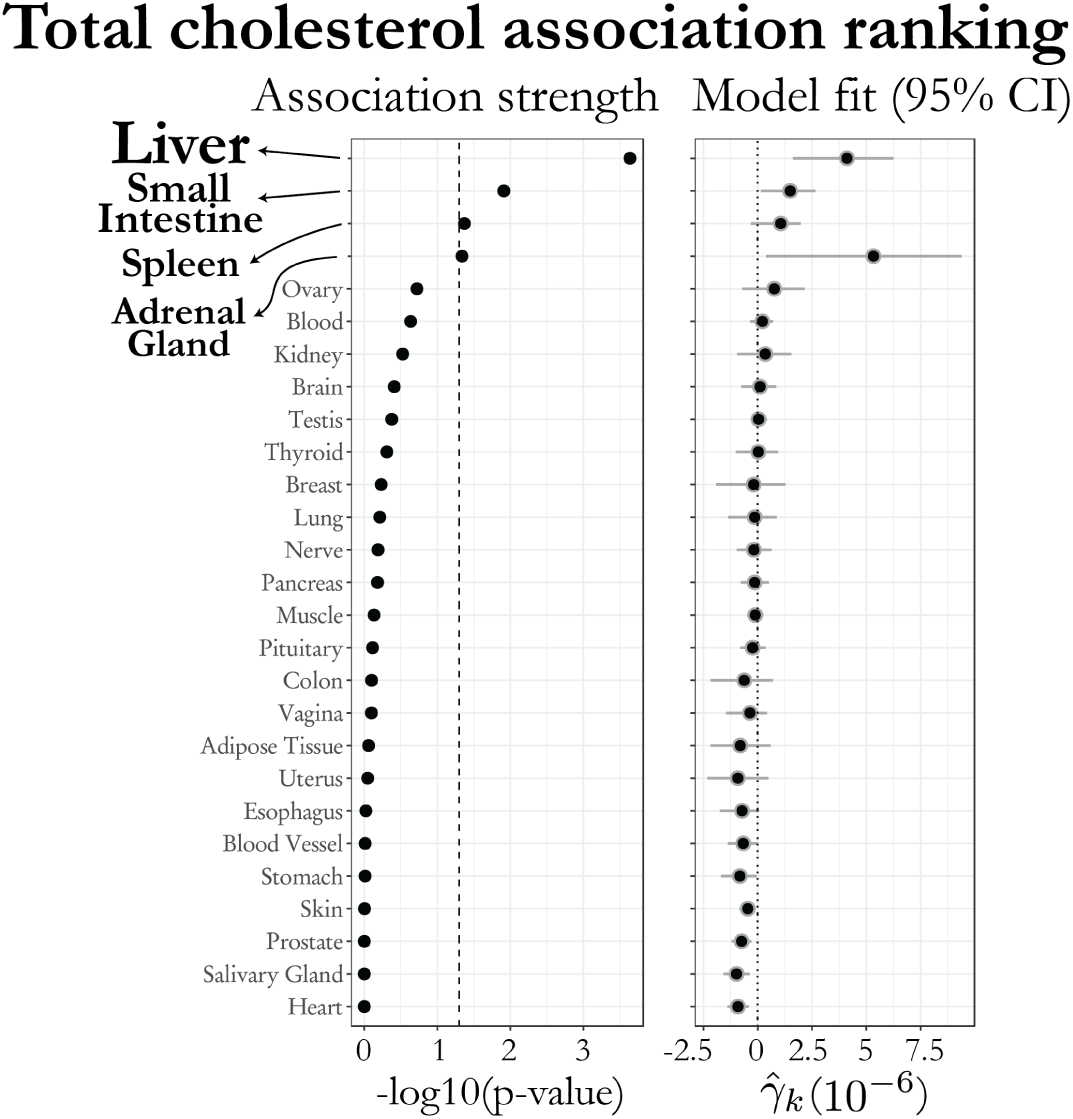
Total cholesterol and GTEx tissue ranking. Left, tissues were ranked by p-value, which represents the strength of association with total cholesterol. Right, corresponding parameter estimates and 95% confidence intervals.

While the spleen is primarily thought of as an immune organ, studies show a clear link between splenectomy and lipid metabolism [13]. While the *p*-value for adrenal gland was identified with a *q*-value of 0.3, the 95% confidence interval show a wide distribution of non-zero parameter estimates of large positive effect. Considering the adrenal gland plays a central role in the production of hormones (many of which are synthesized from cholesterol or even have an effect on cholesterol levels), this association is biologically plausible [42].

We wanted to verify that GWAS effect sizes were enriched for association signal near genes that were specifically expressed in tissues with RolyPoly annotation coefficients significantly greater than zero. First, we calculated LD-informed gene scores, which estimate the variance of GWAS effect sizes from a *cis* window around each gene while accounting for LD (see Material and Methods). Next, we visualized the enrichment of these scores in specifically expressed gene sets using Q-Q plots (Figure 4). To define the set of tissue-specific genes, we sorted normalized expression values for the tissue of interest by decreasing abundance of normalized gene expression and identified the top 20% of genes as the tissue specific gene set. Correspondingly, we refer to the bottom 20% of genes sorted by expression as the control set. We observed clear enrichment of total cholesterol *cis*-GWAS signal within the set of genes that were upregulated in the liver (Figure 4A). As a negative control, we employed the same Q-Q plot approach to determine whether there was GWAS signal around genes specifically expressed in a tissue not found to contribute significantly to total cholesterol. Within specifically expressed genes of the skin tissue (Figure 4B), we did not observe an enrichment of GWAS signal.

**Figure 4:**
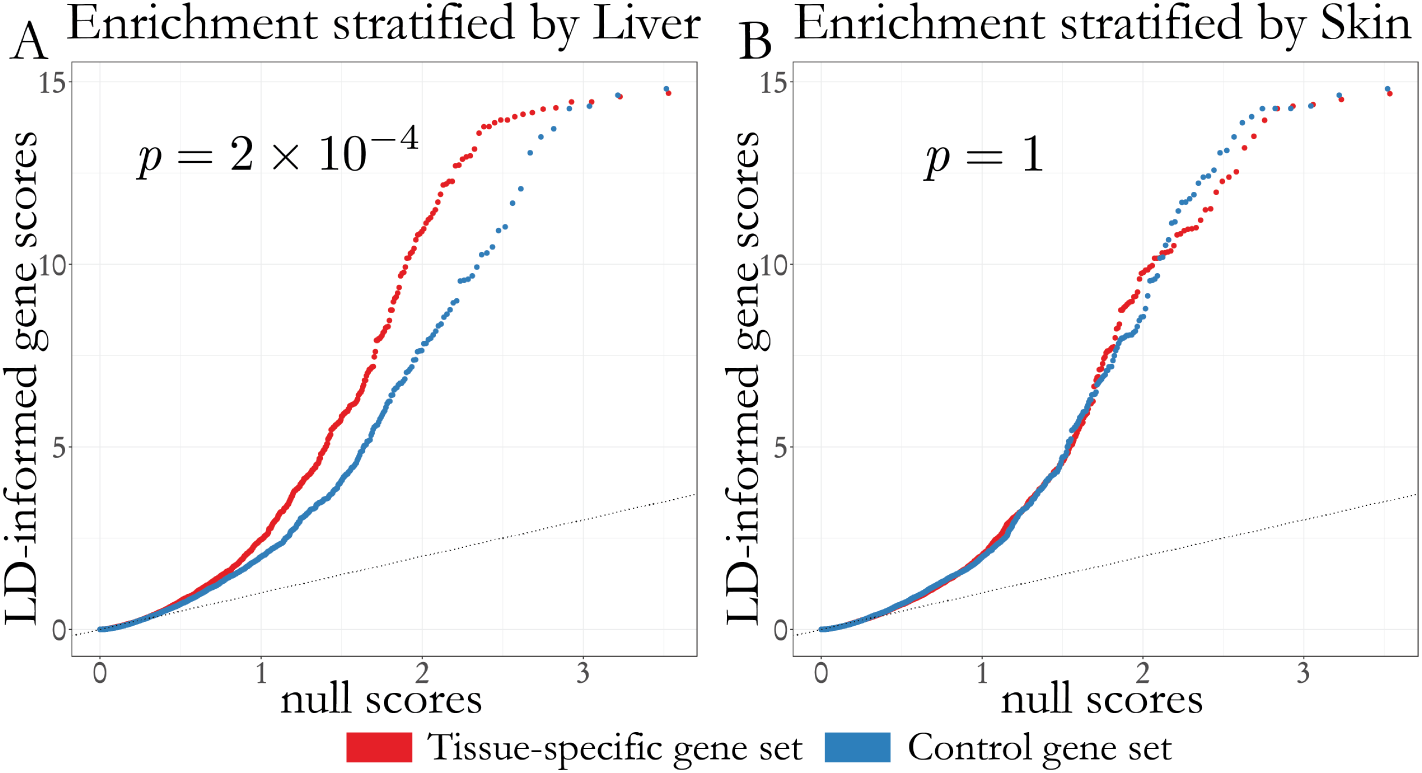
Total cholesterol and GTEx Q-Q plot comparing enrichment of LD-informed gene scores. Both plots show the p-value from RolyPoly for the association between the respective tissue and total cholesterol. A) Q-Q plot comparing enrichment of LD-informed gene scores in genes that are uniquely expressed in the liver. To select gene sets, we sorted genes by their normalized expression in the liver and took the top 20% of genes (red) and the bottom 20% of genes (blue). B) Similar plot, except stratifying gene values by Skin-specific gene expression, a tissue not predicted to have a role in cholesterol regulation.

### Neuropsychiatric diseases and single-cell gene expression

We next analyzed cell types identified from publicly available single-cell expression data from the human brain [7] and several neuropsychiatric traits: age-related cognitive decline, late-onset Alzheimer’s disease, educational attainment, and schizophrenia. In total we used 477 human single cells from which gene expression data were collected. Using a PCA-based clustering approach, the original authors grouped the single cells into 6 primary cell types and two clusters of fetal cortical cells representing quiescent and replicating cell states. For each gene we averaged gene expression counts for all cells within a cell type cluster, thus reducing the noise across single-cell measurements (see Material and Methods). Using our model, we tested the association between each of the traits and 8 clustered cell types (Figure 5).

**Figure 5:**
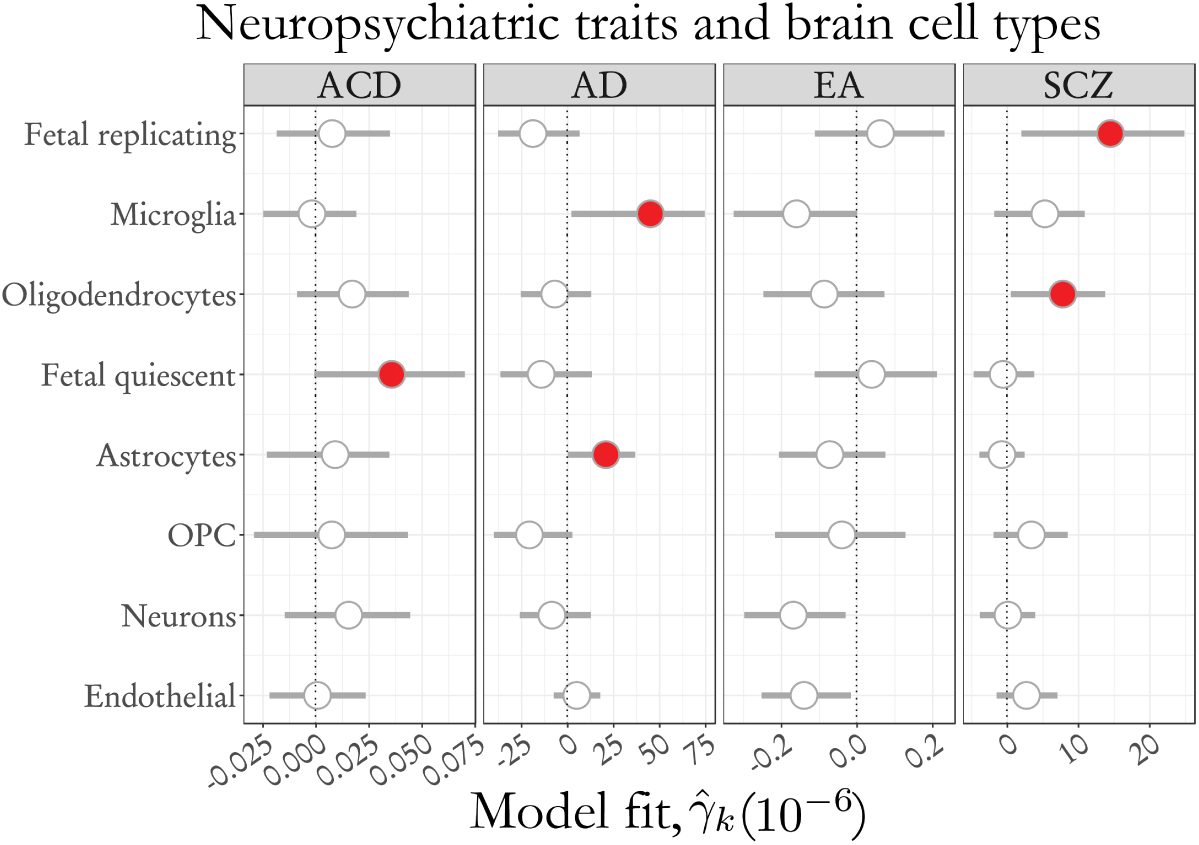
Neuropsychiatric trait associations with single-cell based cell types. Parameter estimates for age-related cognitive decline (ACD), Alzheimer’s disease (AD), Educational attainment (EA) and Schizophrenia (SCZ), and single-cell based cell type clusters from the human brain data set [7]. Range specifies the empirical 95% confidence interval bound. Estimates highlighted in red represent significant associations (p < 0.05).

Age-related cognitive decline (ACD) is a trait characterized by a decline in cognitive capability and decreases in brain volume, both thought to be a normal function of aging. However, evidence suggests that the rate at which cognitive decline occurs is a precursor to late-onset Alzheimer’s disease (AD), hinting at a shared genetic architecture [8]. Thus, we were interested in whether significant overlap of trait-associated cell types existed between the two traits. For ACD, we observed a significant association with fetal quiescent cells (*p* = 0.03), which primarily consists of neurons. Quiescent fetal cells differ from replicating fetal cells in that they have begun to downregulate neuronal growth factors such as EGR1 [7]. On the other hand, we found an association with AD to microglia (*p* = 0.03) and astrocyte (*p* = 0.03) cell types, but no enrichment for fetal neurons (*p* = 0.8). To rule out an association driven by the *APOE* locus we reran RolyPoly while removing a 1 Mb window centered on the *APOE* gene TSS. The significant microglia association persisted (*p* = 0.03) whereas the astrocyte association did not (*p* = 0.1). While the connection between astrocytes and AD is well studied [68], from our analysis this connection appears to be driven by few loci of large effect. Furthermore, there is mounting evidence for a more central role of microglia in AD [18, 56]. However, to our knowledge this is the first human genetics-based enrichment analysis providing evidence for such a connection. Additionally, our results suggest a role for microglia in AD but not ACD. This finding is consistent with recent work demonstrating that while lipid regulation pathways are enriched in GWAS signal for both traits, immune pathways tend to show AD-specific signal [50]. Thus, one could hypothesize microglial involvement during the transition between ACD and AD.

For schizophrenia we found a significant relationship to the oligodendrocyte (*p* = 0.02) and fetal replicating (*p* = 0.01) cell type clusters. The genetic basis of schizophrenia is even less well understood than AD, however there is a significant body of literature studying oligodendrocyte dysfunction and schizophrenia [64, 67]. Moreover, recent genetic association studies have shown an enrichment of schizophrenia GWAS signal within pathways of development [29, 22, 26].

To validate these associations between traits and single-cell cell type clusters, we processed a single-cell data set (see Material and Methods) from mouse brains [71], which included seven major brain cell types that were previously identified. By only utilizing one-to-one human and mouse orthologs, we consider this data set to be an independent pseudo-human brain single-cell data set. Thus, we used this data set to validate our previous findings. We limited our analysis to cell types overlapping in the human and the mouse data set, which included microglia and oligodendrocytes. For AD we replicated the significant association with microglia (*p* = 0.01). Of note, there was a cluster that included astrocytes and ependymal cells, however there was no significant association with this cluster. With schizophrenia there was a suggestive association with the mouse-derived oligodendrocyte cell type cluster (*p* = 0.09). Thus, from our analysis of mouse single-cell data we replicated two of our initial trait and cell type associations. Furthermore, we demonstrate that if human data is not available one could swap in similar mouse data to guide initial analyses.

### RolyPoly gene scores correlate with differentially expressed genes in patients with Alzheimer’s

We were interested in studying whether RolyPoly-inferred model parameters could predict trait-relevant genes from an independent data set. Thus, we downloaded and processed gene expression data from human brain samples of 101 control and 129 Alzheimer’s disease patients from the prefrontal cortex (see Material and Methods and [72]). A total of 9,228 genes were differentially expressed (DE) with a q-value *<* 0.1% (6,324 genes did not meet this threshold). Such a differential expression study represents a data-driven approach to identifying AD-associated genes (independent from GWAS results). Additionally, we used summary statistics from this experiment to test the ability of our model parameters to identify trait-relevant genes.

To establish a baseline, we tested the enrichment of LD-informed gene score estimates within DE genes. These values were computed by taking the variance of GWAS effect sizes within 5kb of a gene and incorporating information about LD (see Material and Methods). We detected only weakly suggestive (*p* = 0.09) enrichment of these values within the set of DE genes compared to genes not found to be significantly expressed.

As a first step to incorporating information from RolyPoly-inferred model parameters, we tested whether genes that were specifically expressed in a RolyPoly-inferred trait-relevant cell type were enriched for larger differential expression test statistics. We identified the top 10% of genes specifically expressed in the microglia cell type (which our model identified as significantly associated with Alzheimer’s disease). Within this set of genes we found a significant enrichment (*p* = 1 × 10^−8^) of positive values of the differential expression test statistic when compared to a control set of genes (right, Figure 6A). We performed a similar analysis with a cell type for which RolyPoly did not find evidence for AD-association. There was no enrichment of DE summary statistic values within the set of genes specifically expressed in fetal quiescent cells (left, Figure 6A).

**Figure 6:**
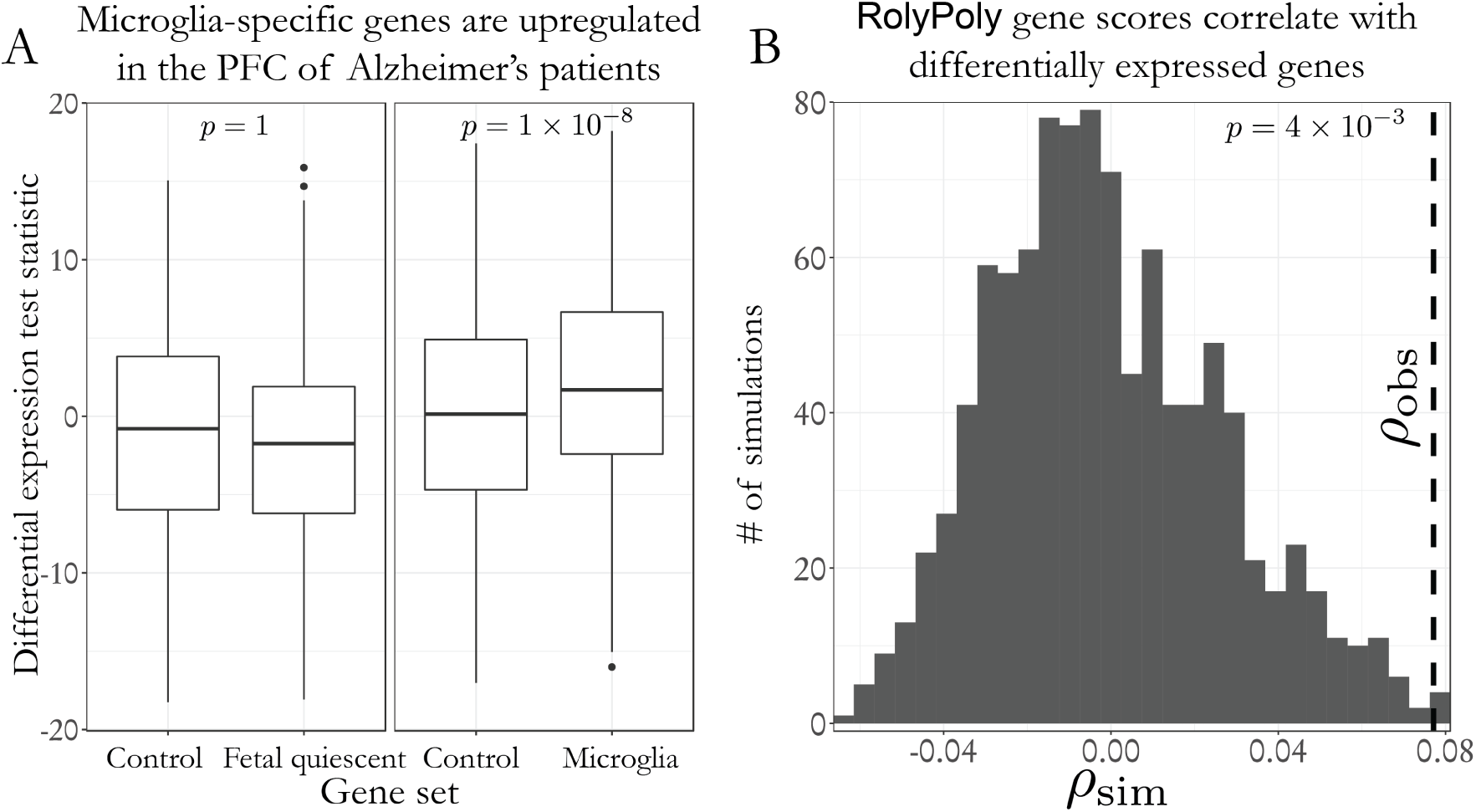
RolyPoly-inferred model parameters predict differentially expressed genes in the prefrontal cortex (PFC) of Alzheimer’s disease patients. A) Differential expression test statistics (a larger value represents genes that are upregulated in the brains of case patients) were significantly larger in the set of genes specifically expressed in the microglia cell type compared with a control gene set (right). We define the set of cell type-specific genes as the top 10% specifically expressed genes. We compared them to the control gene set, which include genes that deviate the least from average gene expression. The differential expression test statistic was not enriched in genes specifically expressed in the fetal quiescent cell type (left). B) Controlling for the effect of correlation between gene expression values of co-regulated genes, we observed an enrichment of 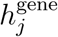 values in differentially expressed genes. The significance of the observed Spearman’s rank correlation coefficient between 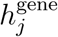 and the differential expression test statistic was evaluated with a null distribution generated from simulations, which account for the gene expression covariance structure (full details of this test can be found in the Material and Methods).

From these observations we reasoned we could rank the trait-relevance of genes based on RolyPoly-inferred parameter estimates,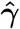, and gene expression. As an example for Alzheimer’s disease, a gene that is specifically expressed in microglia and astrocyte cells would be higher ranked than a housekeeping gene. Thus, we defined the RolyPoly trait-relevance gene score 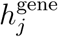 as a linear combination of 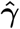 and normalized gene expression values (see Material and Methods). Using the model from the AD-specific panel of Figure 5 and human brain single-cell gene expression we computed estimates of 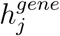. Furthermore, we hypothesized 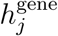 values could predict differentially expressed genes. We found *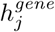* scores were significantly enriched within the differentially expressed genes (*p* = 7 × 10^−18^, Figure S1). However, it is possible that correlations among co-regulated genes could result in uncalibrated *p*-values. Therefore, we designed a test that accounts for the covariance structure between genes (see Material and Methods). Using this test we still identified a significant association (*p* = 4 *×* 10^−3^) between differentially expressed genes and 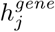 values (see Figure 6B).

For validation, we were interested in replicating our enrichment of 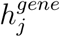 in differentially expressed genes in an independent data set. Sekar et al., performed laser capture microdissection to isolate astrocytes from 10 healthy controls and Alzheimer’s patients, and then identified 227 differentially expressed genes [58]. Of those genes, we predicted RolyPoly gene scores for 150 (the others were excluded because they were not measured in the single-cell expression database). We replicated our previous result and identified a significant (*p* = 1 × 10^−3^) enrichment within differentially expressed genes (Figure S2). We were unable to perform our enrichment test that accounts for gene correlations, because gene expression data was not available for this data set.

Thus, we conclude that from GWAS and gene expression data of healthy individuals our model parameters capture information about the relevance of a gene to a trait based on which cell types express the gene. Still, we cannot discount the possibility that observed enrichments of differential expression test statistics are a result of changes in cell type proportions. However, in such a scenario we would have identified trait-relevant cell types that are increasing or decreasing in proportion and thus would be consistent with our conclusion about RolyPoly parameters.

## Discussion

We described a polygenic model for analyzing single-cell gene expression and GWAS summary statistics. Our results demonstrate that we can identify trait-relevant cell types from complex tissues and prioritize genes for further analysis.

We discuss the following assumptions underlying RolyPoly: i) we focused on *cis*-GWAS effects (as opposed to *trans*), because *cis*-SNPs tend to more consistently have effects, and larger effects, on the regulation of gene expression genome-wide [69, 63, 21, 41, 48]. ii) Our model treats neighboring genes independently even though some may have shared *cis*-SNPs, which could result in correlation among nearby SNP effect sizes. However, we corrected for this effect by performing block bootstrap when computing standard errors and empirical confidence intervals. iii) As this is a joint analysis (we estimate all annotation parameters at the same time), inclusion or exclusion of gene expression data that are causal or correlated with causal cell types can have an effect on inference (i.e., result in different model parameter estimates). However, joint analysis is necessary because analyzing each cell type separately would not control for potential overlap of specifically expressed genes. To mitigate these effects we suggest several approaches. First, we re-analyze a trait GWAS as more data become available. Secondly, we recommend a cautious interpretation of model parameters, which should be guided by domain knowledge. Finally, with highly-correlated annotations, one could carry out an initial round of feature selection before performing standard inference or include regularization (described in Material and Methods). Even with these model assumptions, our results are well supported by known biology, as shown in the analysis of tissues and brain cell types.

To the best of our knowledge, this is the first attempt to connect single-cell gene expression and genome-wide summary statistics from GWAS to identify relevant cell types and genes. While there is evidence linking the immune system and microglia to Alzheimer’s disease [18], we identified for the first time an enrichment of genetic trait-association signal near genes specifically expressed in human microglia. More generally, single-cell technologies represent an opportunity to discover and characterize novel cell types and cell states [52]. Thus, there is a need for methods such as RolyPoly that can prioritize novel cell types for further study that are relevant to human phenotypes. Here, we focused on single-cells clustered into cell types, however there are numerous alternative groupings to examine. For example, during cell stimulation there exists significant cell heterogeneity even within classical marker-defined immune cell type populations [59, 2]. Using RolyPoly one could link these novel sub-populations to autoimmune disease phenotypes. These analyses should only increase as single-cell data become more commonly available.

It is challenging to pinpoint causal genes from GWAS, because correlations among SNP effects due to LD confound the identification of causal variants. Moreover, it is difficult to identify the target gene modulated by a regulatory variant. Statistical methods that integrate GWAS and eQTLs, while accounting for the effects of LD [34, 24], have proven useful. However, the eQTL data may not be specific to the disease-relevant tissue or cell type. To supplement these approaches we suggest using the signature of gene expression and parameters from our model to prioritize genes proximal to significant GWAS variants for further analysis. Consider a region with complex LD structure and significant trait-association signal, ideally one would rely on overlapping eQTL information to identify the causal SNP and gene. But, without knowledge of the causal tissue, GWAS-eQTL overlap with a non-causal tissue could be misleading and complicate the task of collecting relevant eQTL information. Instead, one could use annotation parameter estimates from RolyPoly with tissue or cell type-specific gene expression to calculate *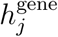* trait-importance values and prioritize genes within the local GWAS region. Additionally, as we have shown, our method can identify significantly associated tissues which one could prioritize for collection of population samples for eQTL analysis.

## Appendix

### Derivation of univariate effect estimates

We follow much of the notation and derivation from [60]. Starting with the definition of the annotation coefficients (recalling that the genotype matrix has been scaled),

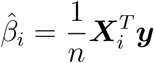

we substitute the GWAS model,

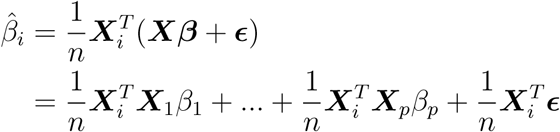

and use the definition of Pearson’s correlation coefficient once again relying on the fact that the genotype matrix have been scaled and centered,

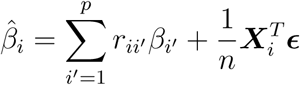

In the Material and Methods section we write the above expression with matrix notation. Others have described a similar relationship between estimated effects, LD, and the true effect sizes [30].

### Derivation of distribution parameters of effect estimates

Here we describe the mean and variance of the estimated SNP effects using our polygenic model. The expected value is computed as follows,

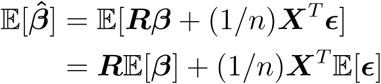

and because we model the genetic and environmental effects with 0 mean normal distributions we conclude that 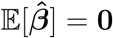 Next,

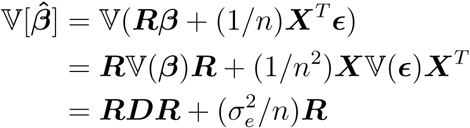

where ***D*** refers to the diagonal matrix of SNP effect size variances and in the second equality we use the fact that ***R*** = ***R***^*T*^. We use these values of the expectation and variance to parameterize the multivariate normal distribution that describes the estimated GWAS effect sizes.

### Derivation of expected SNP variance

Note that the distribution of the squared *ℓ*_2_ norm of a random vector drawn from a mean 0 multivariate normal distribution is the trace of the covariance matrix [37, 40]. Thus, the expected value of the sum of squared SNP effect sizes near gene *j* is given by,

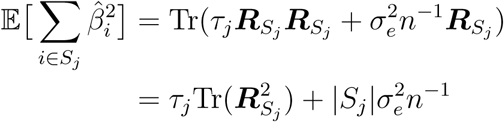

This was derived using the linearity of the trace and recalling that ***R*** is a correlation matrix and hence the diagonal elements are 1. When SNP annotations are included, we model the expected value of the squared marginal SNP effect size. The marginal distribution of the squared SNP effect size around gene *j* is 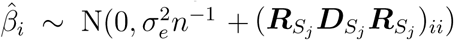. Finally,

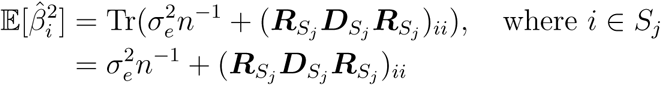

### Relationship to previous work

Rewriting (***RDR***)_*ii*_ as 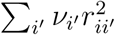, and substituting quantitative feature values with an indicator function that signifies if a SNP is within a discrete annotation class, we arrive at an equation similar to the basic LD Score regression model,

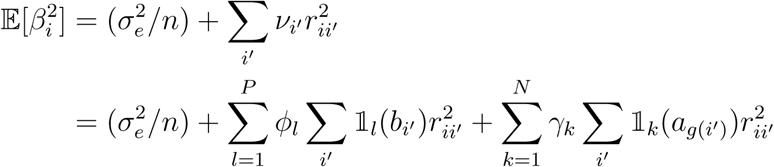

Note that we went from the first to the second line by substituting *v* from equation 3. Although the models share some similarities, our model was derived independently to utilize the full quantitative data from single-cell gene expression assays.

## Supplemental Data

Supplemental Information includes two figures.

## Acknowledgments

We thank Natalie Telis for producing organ images, Ziyue Gao, Naomi Latorraca, and Nasa Sinnott-Armstrong for feedback and discussion, and Anil Raj for early contributions to method development. Support for D.C. was provided by NLM Training Grant Number T15LM007033. This work was supported by NIH grant 1R01HG008140-01A1 and by the Howard Hughes Medical Institute.

## Web Resources

source code repository, https://github.com/dcalderon/RolyPoly

CRAN page, https://cran.r-project.org/package=RolyPoly

